# Eco-toxicity of different agricultural tank-mix adjuvants

**DOI:** 10.1101/2024.09.18.613661

**Authors:** Ivo Roessink, Marie-Claire Boerwinkel, Rick Sijs, Hanne Hoffman, Jan Christof Specker, Daniel Bonn

## Abstract

Adjuvants are often used to improve the efficiency of plant protection products. However, there is concern that these compounds themselves might result in ecotoxicological effects. To investigate this concern, we compare the toxicity of different agricultural tank-mix adjuvants for two standard test organisms, i.e. the water flea *Daphnia magna* and the honey bee *Apis mellifera*. Daphnia trials comprised tests at 1, 5 and 20 times the normal prescribed label dosage. It is found that at 48h, the novel polymer-based adjuvant Squall is significantly less harmful to *D. magna* compared to traditional surfactant or oil-based adjuvants. For *A. mellifera*, we tested topical exposure to label-rate, five and twenty times label-rate. After 96h exposure to polymer- and oil based adjuvants no statistically significant harmful effects were observed. The trisiloxane-based adjuvants, however, did significantly increase bee mortality at higher dose rates, indicating a higher toxicity of these specific compounds.

## Introduction

The growing world population requires agriculture to be as efficient as possible, and without pesticides this is difficult. The US Academy of Science report^1^ shows that the use of efficient pesticides has more than doubled the productivity in US agriculture, and that agriculture without pesticides is unthinkable nowadays. Although efficiency is key, it is reported that over 50% of the average crop protection spray reach a destination other than their target species. ^2^ Other major parts of pesticides are lost to, groundwater, surface water, or adjacent soils due to wind drift, surface run-off, or soil drainage. ^3–5^ Bounce or run-off of the treated surface is the main loss mode for agricultural pesticide sprays, significantly reducing the efficiency of deposition and retention of droplets on plant surfaces.^6–8^ Consequently, adjuvants are often added to the tank mix to enhance droplet spreading and sticking on target surfaces, as well as penetration into leaves and barriers for a more active ingredient uptake.^9,10^ Usually, adjuvants are much less expensive than formulated active ingredients and can reduce the active ingredient dose needed to reach efficacy by an order of magnitude or more.^11,12^ While active ingredients of plant protection products are strictly checked for environmental impact, the same does not always apply to adjuvants; regulations vary from country to country, but in most cases the requirements are much less strict than for pesticides. That this can be an omission, is shown by the example of the herbicidal active ingredient Glyphosate that in a formulation with adjuvants (RoundUp) is more harmful than in its pure state.^13^ Spray tank adjuvants have also been shown to be potentially toxic to bacteria,^14^ cyanobacteria,^15^ algae,^16^ snails, ^17^ *Daphnia pulex* ^18^ and insects,^19^ but also invertebrates, ^20,21^ fish,^19,22,23^ and amphibians^24^ can be impacted.

In this article, we investigate the toxicity of a few commonly used adjuvants for the two standard test organisms the water flea *Daphnia magna* and the honey bee *Apis mellifera. D. magna* represents aquatic invertebrates that can be indirectly exposed to any spray drift reaching edge-of-field surface waters, whereas *A. mellifera* can be directly exposed to these compounds via over-spray or contact via treated plant surfaces. There are many agricultural adjuvants on the global market, and for this first study we have limited ourselves to a number of adjuvants that are commonly used in Western Europe, still spanning a range of active ingredients that is as broad as possible. Further studies are undoubtedly needed to include also other (types of) adjuvants. The adjuvants used and their chemical characteristics are listed in table 1.

**Table 1:** The different Adjuvants used in the toxicity tests with Daphnia Magna and Apis mellifera, and some of their chemical and physico-chemical properties.

| Adjuvant | Chemical composition | Group | Manufacturer | Surface tension (mN/m) | Advised concentration |
| --- | --- | --- | --- | --- | --- |
| Prolong XP | Solvent Nafta (Petroleum) | Spreader | InterAgro | 32.5 | 0.15% |
| Driftstop | Organosilicone surfactant synthetic polymer | Spreader<br>Sticker<br>anti-drift | Nufarm | 20.9 | 0.2% |
| Certain | alkoxylated alcohol | Spreader | Modify | 25.8 | 0.1% |
| Zipper | Polyether modified polytrisiloxane | Super wetter | Modify | 21.8 | 0.1% |
| Wetcit Neo | Alcohol, C12-14, ethoxylated, sulfates, sodium salts | Spreader | Oro Agri | 33.5 | 0.3 % |
| Squall | Poly-ethylene Oxide | Spreader<br>anti-drift | GreenA BV | 30.3 | 0.5 % |
| Silwet Gold | Heptamethyl-trisiloxane | Super wetter | Arysta | 20.6 | 0.075 % |
| Breakthru S240 | Trisiloxane | Super wetter | Evonik | 21.6 | 0.1 % |
| NuFilm P | Pinene (polyterpenes) polymers, petrolatum | Spreader | Miller chemical | 37.2 | 0.15 % |
| Hi-Wett | Polysiloxane polyether copolymer | Super wetter | Nufarm | 21.5 | 0.2 % |

**Table 2:** EC_50_ values (mg L ^-1^) with their corresponding standard error (SE) and 95 % confidence interval (CI) over 48 h for different adjuvants to *D. magna*.

| Adjuvant | EC<sub>50</sub> (mg L<sup>-1</sup>) | SE | 95 % CI |
| --- | --- | --- | --- |
| NuFilm P | 0.59 | 0.07 | 0.46 - 0.72 |
| Hi-Wett | 2.48 | 0.44 | 1.58 - 3.37 |
| BreakThru S240 | 2.48 | 0.44 | 1.58 - 3.37 |
| SilWet Gold | 6.10 | 0.95 | 4.20 - 8.01 |

## Materials and methods

### Daphnia magna

*Daphnia magna* is a standard organism often used in assessing chemical toxicity, following OECD guideline 202.^25^ Our study followed the OECD 202 test set-up to determine the 50% effect concentration (EC_50_) leading to immobilization for each adjuvant after 48h exposure to different test concentrations. Daphnids were obtained from the in-house daphnid culture of the laboratory and were not older than 24h at the start of the test. To examine their fitness, parallel toxicity tests were performed with the reference toxicant potassium-dichromate. Daphnid survival indeed fell within the approved range indicating the population was fit-for-purpose. For each test solution, five daphnids were placed in a 10mL glass vial filled with the diluted substances or control medium. The vials were placed in a climate chamber with a 12h:12h light-dark cycle at a temperature of 21^*°*^C for 48 hours. No feed was added during the experiment.

Two series of toxicity tests were performed. The first series was a broad range finding experiment, in which a broad range of substance dilutions was used to find approximate toxicity values. The second series was a precision toxicity test, to fine tune the toxicity for each of the substances. During range finding, four dilutions were tested with a sample size of n=2. During precision testing, five or six dilutions were tested at 1, 5 and 20 times the normal prescribed label dosage, with a samples size of n=4. Control vials containing medium only were used during both series. Effect values are expressed as EC50.

### Honey Bees

Acute contact exposure tests were performed with honey bees according to the OECD guideline 214 (OECD, 1998b). Bees were collected from queen-right colonies at the Sinderhoeve field station, the outdoor test facility of Wageningen Environmental Research. Tests were performed in triplicate (n=3) and each replicate held 10 individuals. Replicates were populated with individuals originating from one colony and three colonies were used to populate the three replicates per treatment. As these compounds were not tested before we prolonged the test until 96 hours after dosing. In an acute contact test (OECD, 1998b), the substance to be tested is applied as a 1–2 *µ*L droplet on the dorsal thorax of the bee. In our tests, water was used as a carrier liquid and the addition of the adjuvants ensured the droplet flowed evenly over the thorax, thus ensuring adequate exposure. Since the contact between the adjuvants and the bees can be direct, we directly tested the different adjuvants at 1, 5 and 20 times the normal prescribed label dosage of the adjuvants. Since water evaporated rapidly from small spray droplets, there is a risk of exposure to a higher concentration than the label dosage; hence the inclusion of 5 and 20 times the label dosage. After dosing the bees were put in ventilated plastic containers and were fed *ad libitum* with a uncontaminated 50%:50% sugar:water solution. Mortality was recorded at the designated time points for the remainder of the observation period.

Table 1 lists the selected adjuvants with their ingredients. The medium in which these substances were diluted is the ADaM medium for Daphnia tests, consisting of: CaCl2.2H2O, 2.0mM; MgSO4.7H2O, 0.5 mM; NaHCO3, 0.7 mM; KCL, 0.07 mM. This is the only deviation from the guideline, where the standard ISO medium is proposed.

### Data Analysis - *D. Magna*

Data analysis was performed with R^26^ in RStudio, Version 2022.7.2.576, ^27^. The concentration-response relationships were calculated and fitted with the aid of the drc package^28^. For each adjuvant, different models were tested such as log-logistics models with 2, 3, or 4 parameters. Based on the akaike information criterion (AIC), the model with the lowest AIC was chosen. Furthermore, after selecting the model that lead to the best fit to the data according to the AIC, an anova test was run to further compare the model fit between the models. Using the “ED()” function of the drc package with the interval set to “delta”, EC_50_ values were calculated with their standard error and corresponding 95 % confidence interval ^28^.

## Results

### Daphnia magna

An example of two dose response curves is shown in figure 1. The EC50 is the value at which the dose-response curve reaches 50%.

**Figure 1:**
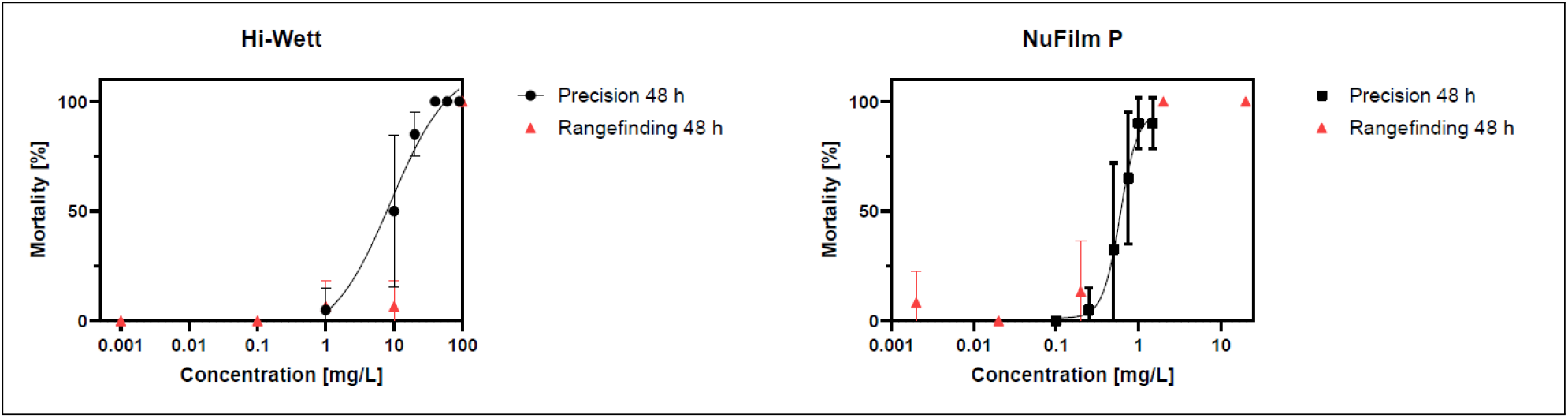
Example of the dose-response curves for Hi-Wett and NuFilm P.

There are large differences in the EC50 values between the different adjuvants, see figure 2. These values are obtained from the dose-response curves that were obtained from the daphnia magna toxicity test. An example of two dose-response curves is given in Appendix A. Squall is by far the least harmful adjuvant with an EC50 of 3909 ul/L. The different groups of chemical compositions are clearly reflected in the results. The (synthetic) polymers such as Driftstop and Squall show the lowest mortality as reflected by the highest EC_50_(48h) values. In contrast, all the surfactant- or oil-based adjuvants give a much high mortality rate. The different types of trisiloxanes give an EC50 of between 6 and 27 uL/L and the most toxic chemical component appears to be petroleum, with an EC50 of 9.5 uL/L for Prolong XP and 0.6 uL/L for NuFilm P. The surface tension appears not to be strongly correlated with the EC50: from our data it follows that the lowest surface tension does not always produce the largest mortality (lowest EC50).

**Figure 2:**
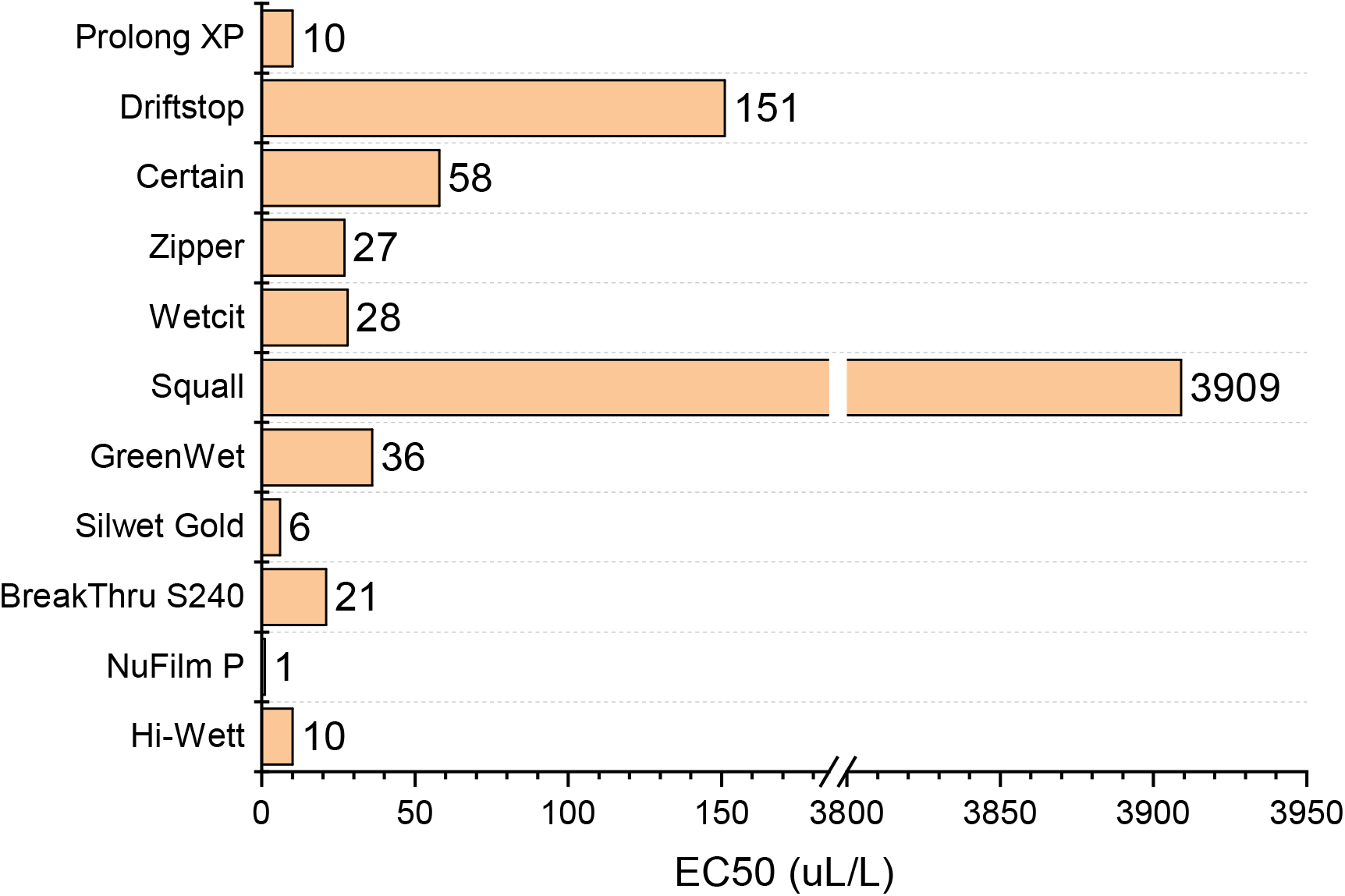
The 48h-EC50 values for the toxicity of different tank-mix adjuvants to *Daphnia magna*

**Figure 3:**
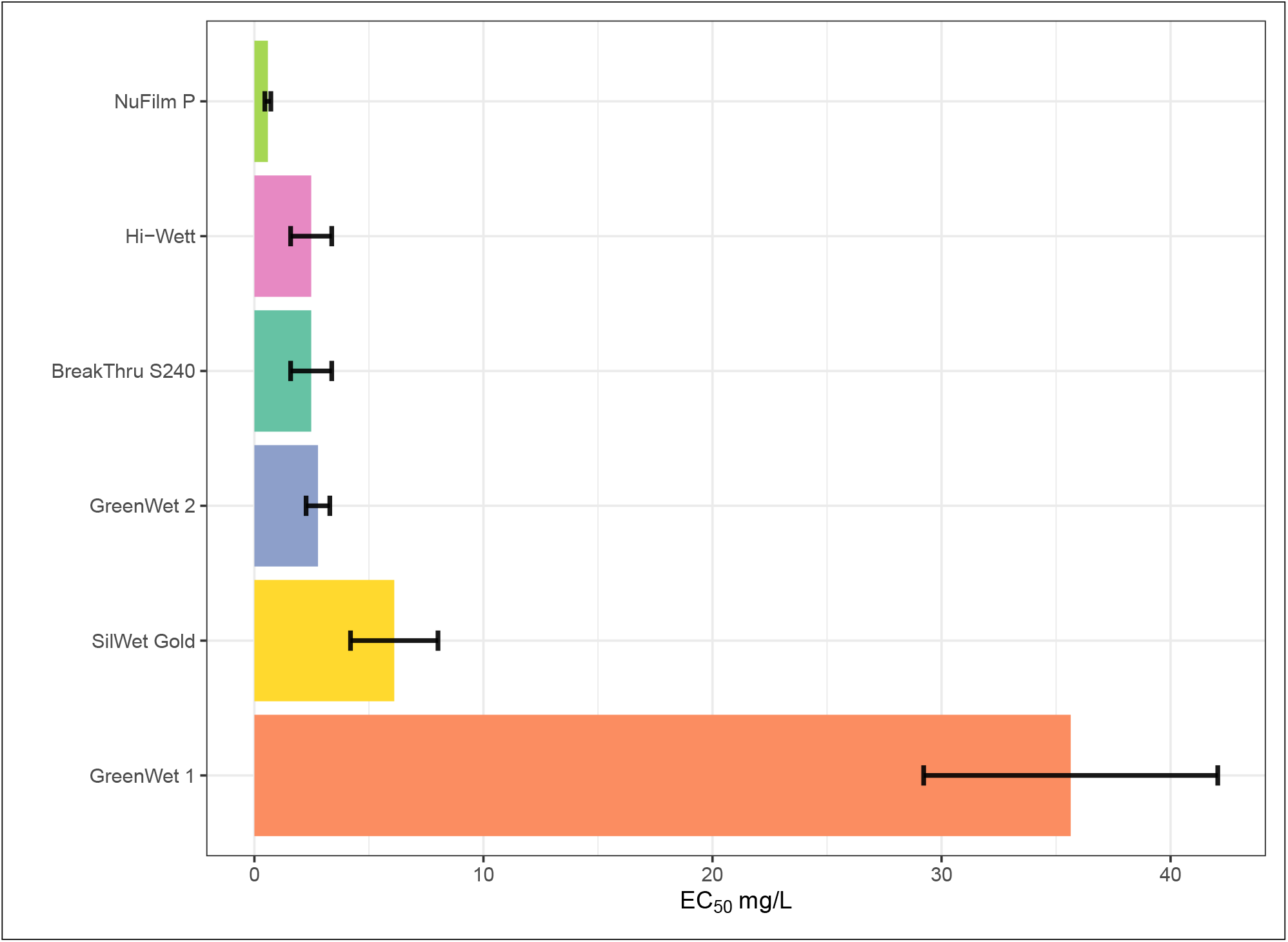
Acute toxicity over 48 h of different tank-mix adjuvants to *D. Magna* expressed as EC_50_ values in mg L ^-1^. Error bars indicate the 95 % confidence interval.

### Honey Bees

Figure 4 shows the results for the honey bee survival. Honey bee survival after 96hrs is rather high in all tests performed, except when testing at 20 times the label dosage. In the latter tests, the three trisiloxane-based adjuvants (Zipper, BreakThru and Hi-Wett) give a significantly higher bee mortality than the other adjuvants.

**Figure 4:**
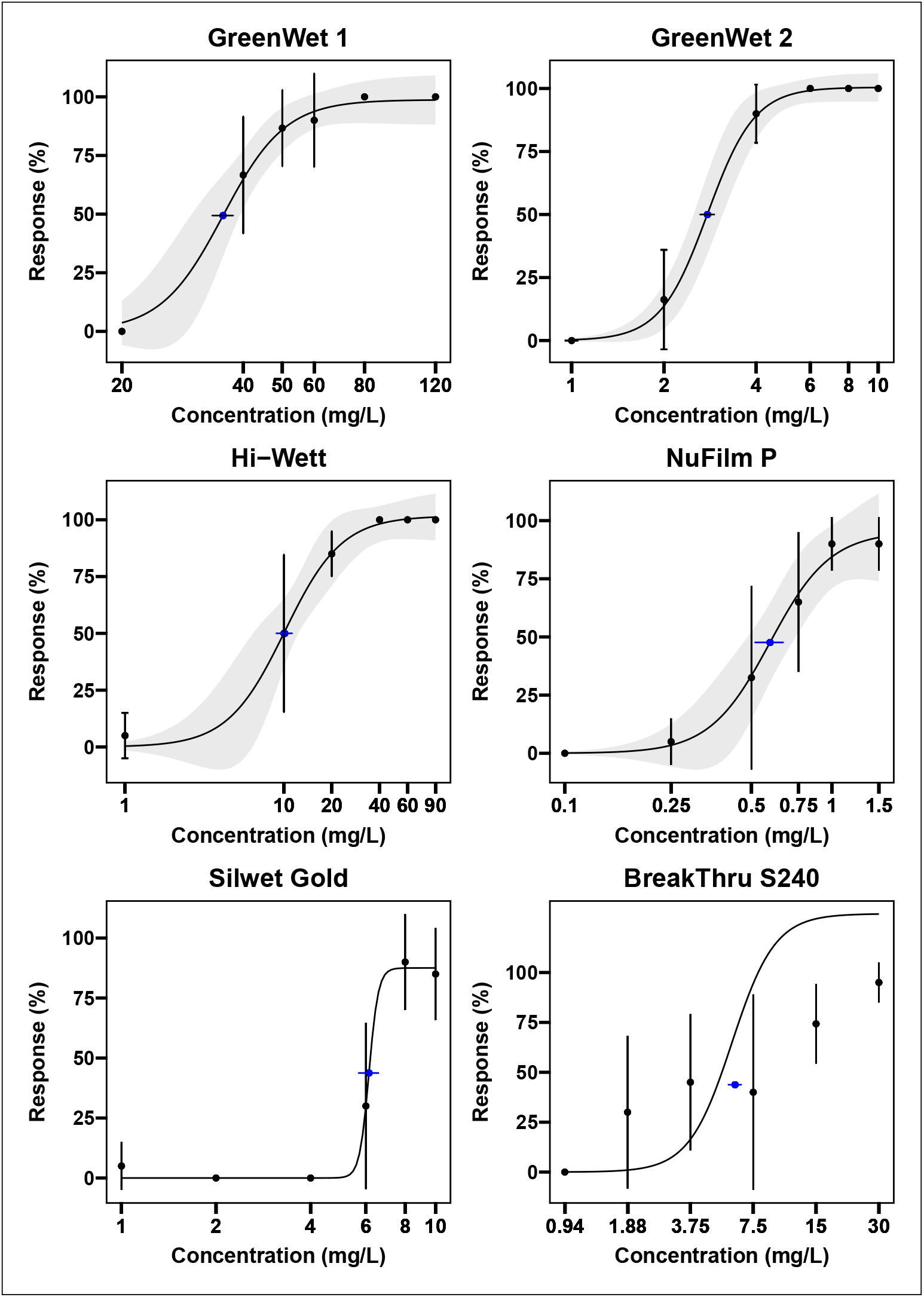
Concentration-response curves for different adjuvants over 48 h exposure. Black points indicate the observed average response with their standard deviation indicated as vertical bars around it. The blue point marks the EC_50_ with its standard deviation shown as horizontal bars. The fitted curve, based on a three-parameterr log-logistic model, represents the concentration-response relationship. The gray shading around the fitted curve shows the 95 % confidence interval.

**Figure 5:**
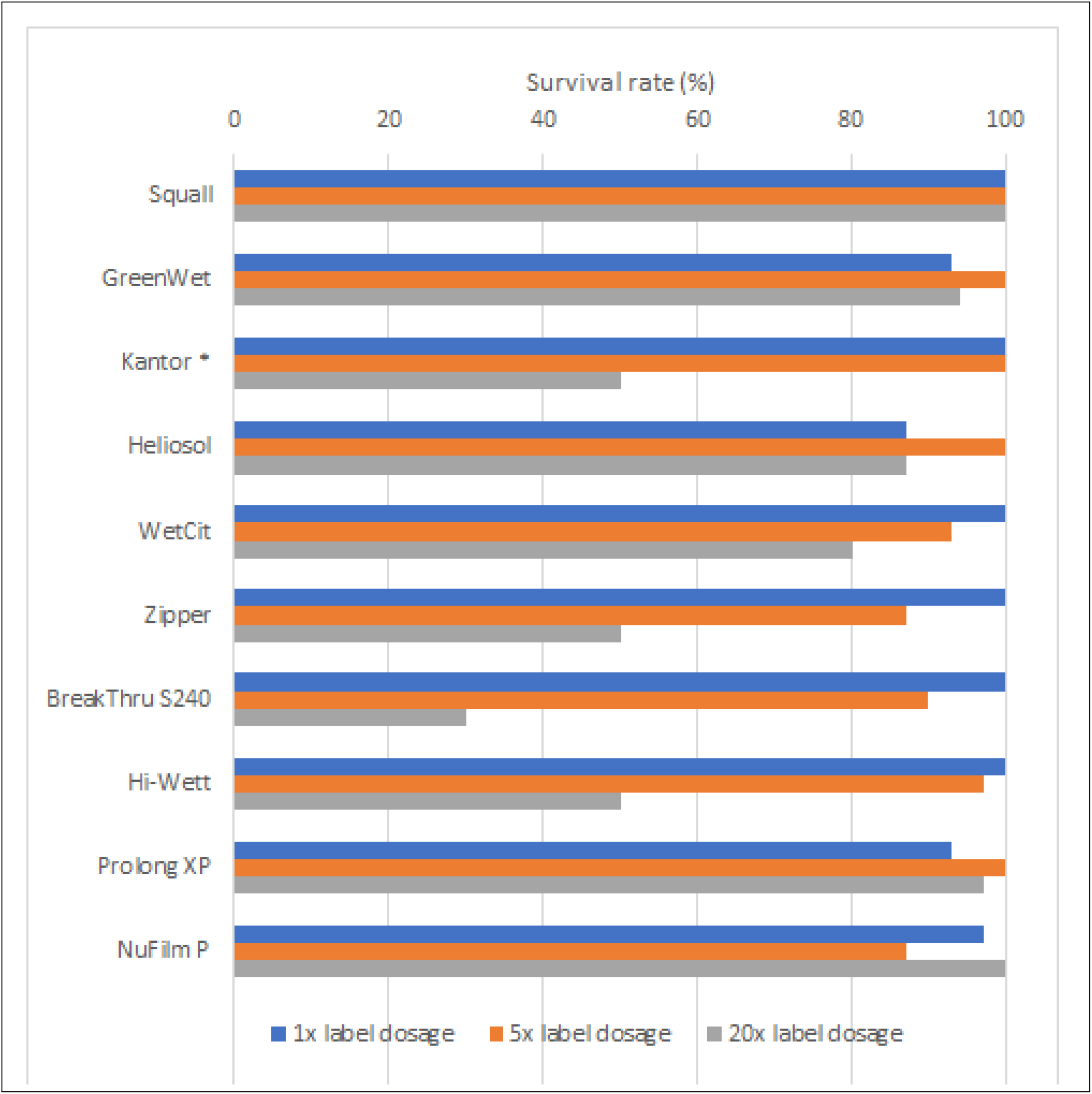
Bee survival after 96 hrs at 1, 5 and 20 times the label dosage

## Conclusion

The observation that the tested adjuvants are harmless for bees at label and 5 times label dosing is reassuring, in the light of the observation made earlier that a study on Glyphosate shows that in a formulation with adjuvants (RoundUp) it is more harmful to bees than pure Glyphosate. For *D. magna*, one of the standard test organisms representative for aquatic invertebrates, on the other hand, only the polymer-based adjuvants appear to be harmless, with Squall still being more than an order of magnitude better (less harmful) than Driftstop. In contrast, the oil- and surfactant based adjuvants appear to be harmful to very harmful for aquatic life. This risk is taken into account when adjuvants are part of the commercially available formulated product and are integrally tested in the regulatory process. However, when sold separately as tank additives adjuvants are not part of the regulatory process and might be a reason for concern. We therefore suggest that these compounds are also subject to a screening to asses potentially occurring environmental concentrations and thereby impact analogue to the active ingredients.

Furthermore, the authors recommend that future studies should investigate chronic effects on *D. magna, A. mellifera*, or other organisms to better understand potential effects on individual organisms but also on the population level due to long term exposure; the tests here were performed for only a few days and the adjuvants may persist for longer than that and/or be used in multiple applications. These tests could also be supported by biodegradation experiments of different environmental compartments such as soil, water, or water-sediment. A better understanding of adjuvant’s persistency and mobility, their fate and behaviour in the environment, would help to better predict their long time effects and transport.

## Acknowledgments

This work is part of an Industrial Doctorate contract at the University of Amsterdam that is supported by the Netherlands Organization for Scientific Research (NWO) under project-number NWA.ID.17.016.

